# Precise, Specific, and Sensitive *De Novo* Antibody Design Across Multiple Cases

**DOI:** 10.1101/2025.03.09.642274

**Authors:** Yewon Bang, Hyeonjin Cha, Kyesoo Cho, Jeonghyeon Gu, Daehyeon Gwak, Young-Hyun Han, Mirim Hong, Suyeon Kang, Sohee Kwon, Changsoo Lee, Dohoon Lee, Myeong Sup Lee, Sangchoon Lee, Su Jung Lee, Jungsub Lim, Jin Young Maeng, Jinsung Noh, Hyunjeong Oh, Soyeon Oh, Seongchan Park, Taeyong Park, Seongok Ryu, Deok Hyang Sa, Chaok Seok, Moo Young Song, Jonghun Won, Hyeonuk Woo, Jinsol Yang, Min Ji Yoon

## Abstract

The precision design of antibodies, which naturally recognize diverse molecules through six variable loops, remains a critical challenge in therapeutic molecule discovery. In this study, we demonstrate that precise, sensitive, and specific antibody design can be achieved without prior antibody information across eight distinct target proteins. For each target, binders were identified from a yeast display scFv library of approximately 10^6^ sequences, constructed by combining 10^2^ designed light chain sequences with 10^4^ designed heavy chain sequences. Binders with varying binding strengths were identified for all eight targets, including a case where no experimentally resolved target protein structure was available, demonstrating the highest level of precision compared to previous *de novo* antibody design reports. To further validate the designed antibodies, they were characterized in the IgG format for five distinct targets. The antibodies exhibited favorable developability properties, positioning them as promising hit or lead candidates. Notably, for one target in particular, the IgG-formatted antibodies exhibited affinity, activity, and developability, comparable to a commercial antibody, highlighting the sensitivity of the design method. Furthermore, binders capable of distinguishing closely related protein subtypes or mutants were identified, demonstrating that the method can achieve high molecular specificity. Cryo-EM analysis experimentally validated the design’s accuracy by confirming that the key binding interactions were precisely as designed. These findings underscore the effectiveness of precision molecular design based on atomic-accuracy structure prediction. This study establishes computational antibody design as a viable approach for generating therapeutic molecules with tailored properties, with promising potential for achieving the efficacy and safety required for successful therapeutics.

## 1 Introduction

The precise *de novo* design of antibodies is a highly valuable approach in therapeutic antibody discovery. Unlike conventional methods that rely on existing immune repertoires, *in silico* antibody design enables the generation of novel binders with tailored molecular properties. A key advantage of this approach is its ability to precisely target specific epitopes for controlled functional regulation of the target protein, providing a level of precision unattainable with traditional methods. Additionally, it allows for the targeting of challenging epitopes that are difficult to address using conventional approaches due to experimental limitations, broadening its potential application to a diverse range of disease targets. By translating prior knowledge of molecular mechanisms of action into molecular properties, this strategy enables the precise design of molecules with optimized therapeutic efficacy and reduced toxicity by enhancing target regulation while minimizing both on-target and offtarget interactions.

Compared to *in silico* generation of protein binders that interact through secondary structure elements—such as *α*-helices or *β*-strands—[e.g., RFdiffusion [1], AlphaProteo [2]], antibody generation presents a greater challenge due to the intrinsic structural variability of complementarity-determining region (CDR) loops. This complexity arises from the limited availability of atomic-resolution structural data for loop-mediated protein–protein interactions compared to those involving secondary structure elements. The high conformational flexibility of CDR loops complicates both structure prediction and affinity estimation, further increasing the design difficulty. As a result, achieving precise control over target binding mediated by CDR loops remains a significant challenge in *de novo* antibody design.

Notably, the Baker Lab [3] and Nabla Bio [4] have recently reported successful *de novo* design of single-domain antibodies that bind their targets via three loops, marking a significant advancement in the field. Additionally, the same groups have reported a few noteworthy *de novo* two-domain antibody designs that engage their targets through six loops. However, the limited number of successful cases, lower success rates, and relatively weaker affinities compared to previous *de novo* binders utilizing secondary structure elements highlight the need for further improvements in this field.

In this report, we present a significant advancement in *de novo* antibody design, demonstrating its *in silico* performance on benchmark test sets compared to existing design methods, as well as its application to eight therapeutic targets followed by *in vitro* validation. The computationally designed antibodies were validated in the single-chain variable fragment (scFv) format, exhibiting precise and specific binding to their target proteins. We evaluated the biophysical properties of Immunoglobulin G (IgG)-formatted antibodies for five different targets. Antibodies produced in the IgG format exhibited biophysical properties that would be considered favorable for potential hit or lead compounds, demonstrating a high level of binding affinity and developability. Especially, for one target, the designed antibodies also demonstrated binding affinity and functional properties comparable to or superior to those of a commercial therapeutic antibody, underscoring the sensitivity of our *de novo* design.

These findings present new challenges in *in silico* biomolecular design, particularly in targeting complex epitopes that are difficult to address experimentally and in designing molecules with environment-sensitive properties, such as pH dependence, as the next stage of advancement. Overcoming these challenges will expand the range of accessible disease targets and facilitate the development of therapeutic molecules with optimized molecular properties, enhancing efficacy and reducing toxicity.

## 2 Results and Discussion

### 2.1 *In silico* evaluation of *de novo* antibody design

The *in silico* evaluation of antibody design was conducted in two parts: (1) assessment of antibody generation performance and (2) evaluation of binder/non-binder discrimination performance.

***In silico* assessment of *de novo* antibody generation performance** We evaluate our *de novo* antibody generative method, GaluxDesign (v2, v2.1, and v3), which builds upon and improves the previously reported antibody loop design method based on the Galux structure prediction model [5]. Its performance is compared to RFantibody [3], which utilizes the RFdiffusion model finetuned for antibody backbone generation and ProteinMPNN for side-chain design, and dyMEAN [6]. Other recently reported methods, such as IgGM [7], could not be benchmarked due to licensing restrictions. Detailed inference protocols for the compared models are provided in Methods (Subsection 4.2).

Here, the *de novo* antibody generation task is defined as generating new variable fragment (Fv) structures and corresponding variable heavy chain (VH) and variable light chain (VL) sequences based on a given set of amino acid residues that define the epitope of the target protein. To ensure an objective benchmark test, a non-redundant dataset of 32 antibody–protein complexes was curated from the Protein Data Bank (PDB), comprising moderate-sized target proteins with high-resolution experimental structures (Methods, Subsection 4.1). Structure quality of the generated antibodies (Figure 1A) and the reproducibility of reference antibody orientation (Figure 1B) were evaluated for 50 *de novo* designed antibodies for each target protein, given five epitope residues closest to the reference antibody in the PDB structure.

**Figure 1.**
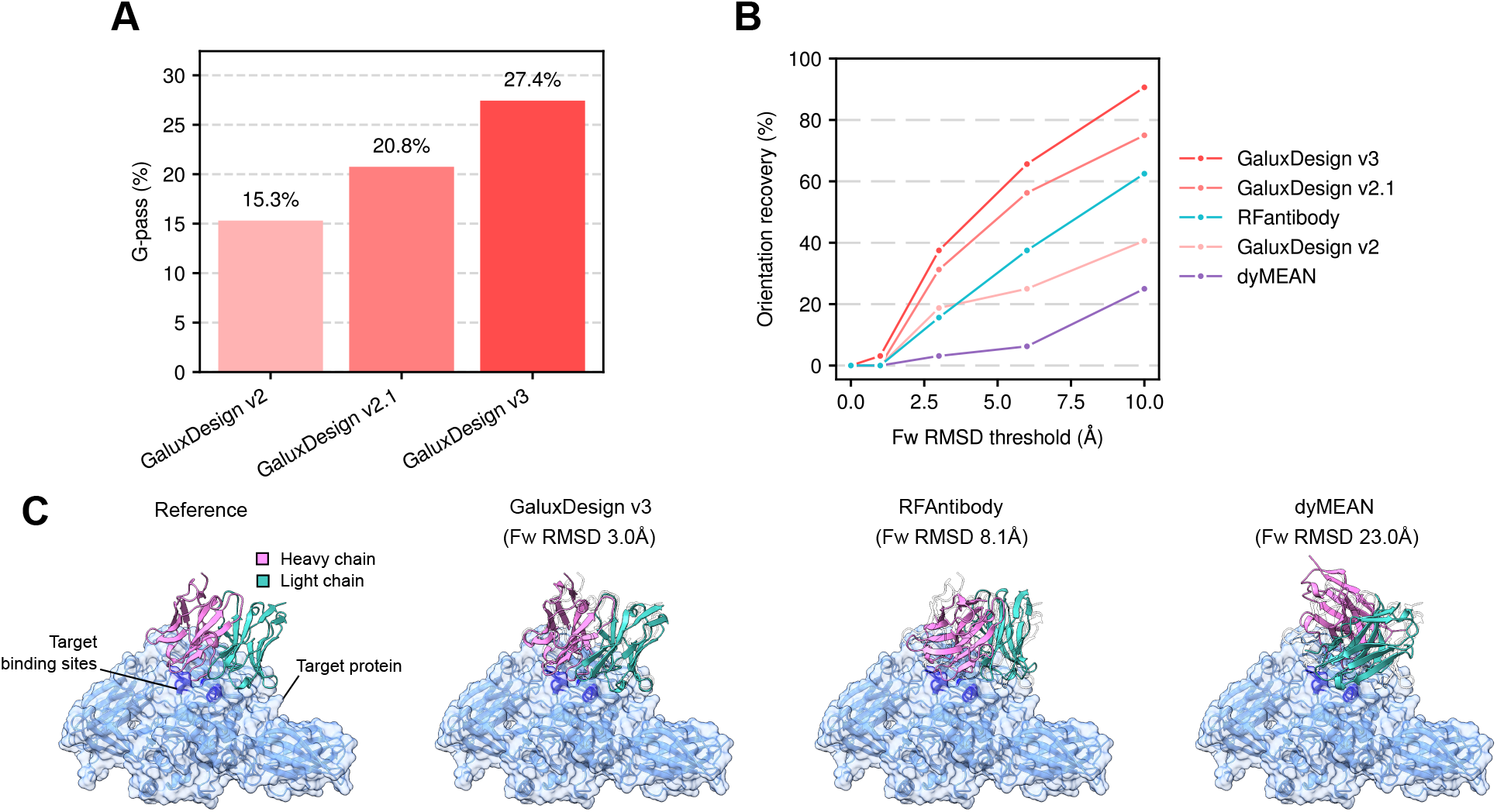
*In silico* benchmark results of *de novo* antibody generation. (A) G-pass rates of the GaluxDesign models, (B) orientation recovery rates across varying Fw RMSD thresholds, and (C) representative *de novo* antibody design results for one of the largest target proteins in the benchmark set (Transglutaminase 3, PDB entry 8OXW). For each method, the design with the lowest Fw RMSD among 50 generated antibodies is shown. Transparent antibody structures indicate the reference antibody. Fw RMSD denotes Root-MeanSquare Deviation of Framework residues after aligning target protein structures.

Figure 1A shows the G-pass rates of the *de novo* designed antibodies across the three versions of GaluxDesign, demonstrating a steady improvement from 15% to 21% and 27%. The G-pass rate, initially introduced for the *in silico* evaluation of antibody loop design [5] with a given antibody orientation, assesses the effectiveness of each antibody design by evaluating the confidence of the predicted structure and the consistency between the designed structure and the predicted structure, as estimated by Galux structure prediction model v2.

For *de novo* antibody design, the G-pass rate of GaluxDesign v2 decreased to 15% from 26%, which was observed in a simpler loop re-design task that leveraged antibody orientation information from the PDB structure of a reference antibody complexed with the target [5]. This result highlights the inherent challenge of *de novo* design without prior antibody information compared to designs incorporating such information, although a direct comparison is challenging due to differences in tasks and benchmark sets. The consistent improvement across model updates underscores the synergy between the enhanced Galux structure prediction model and the design strategy built upon it. Notably, RFantibody and dyMEAN failed to achieve any measurable G-pass rates on the same benchmark set, though this comparison may be biased due to the use of the Galux structure prediction model.

Next, we examined how well the set of 50 antibodies designed by each method covered the space of the reference antibody, given the five epitope residues on the target protein closest to the reference antibody in the available PDB structure. Although higher coverage does not necessarily indicate a better design method, this measure serves as a practical proxy given the high cost of experimental validation. To evaluate this, we compared the proportion of targets for which at least one designed antibody reproduced the reference antibody–protein orientation within a specified threshold, as measured by the Root Mean Square Deviation of Framework residues (Fw RMSD) after structural alignment of the target protein (hereafter referred to as orientation recovery). Figure 1B shows that GaluxDesign v3 and v2.1 outperformed the other methods, while RFantibody exhibited measurable recovery, particularly at larger orientation thresholds. A representative example is shown in Figure 1C, where GaluxDesign v3 recovered the reference antibody orientation with an Fw RMSD of 3 Å, compared to 8 Å for RFantibody.

Altogether, GaluxDesign, built upon accurate atomic-level structure prediction, demonstrates excellent generative performance in *de novo* antibody design across diverse epitopes. This is evidenced by higher G-pass rates, reflecting the effective generation of precise loop-mediated protein–protein interactions, and higher orientation recovery, indicating robust coverage of antibody space.

#### *In silico* assessment of antibody binder/nonbinder discrimination performance

Although the generative model inherently evaluates structural constraints when designing antibodies, a more precise scoring function is required to select the most promising candidates by performing higher-resolution assessments of predicted complex structures and binding affinity. Given the vast number of possible amino acid combinations across numerous variable sites, the importance of an accurate scoring function is even more pronounced in *de novo* design. To address this, we developed an enhanced scoring function that improves upon our previous work [5] by better capturing the fundamental principles of protein–protein binding, leading to improved scoring performance.

To evaluate the performance of the newly developed scoring function, we assessed its ability to discriminate between binder and non-binder sequences across ten target proteins from three different studies [8, 9, 10], each consisting of hundreds to hundreds of thousands of sequences. Detailed information about the datasets used in this evaluation can be found in Methods (Subsection 4.1). For datasets containing tens of thousands of sequences or more, a subsample of approximately 1,100 to 1,200 sequences was utilized for evaluation to facilitate faster computation.

As shown in Figure 2, Galux v3 scoring func-tion demonstrated excellent discriminative power, achieving an Area Under Receiver Operating Characteristic Curve (AUROC) of more than 0.8 in 7 out of 10 datasets, with an average AUROC of 0.80. Specifically, in the HER2–Trastuzumab, 5A12–Ang2, and 5A12–VEGF antibody libraries containing the largest number of binder/non-binder data, the method achieved AUROCs of 0.80, 0.86, and 0.89, respectively, reflecting its strong performance. These results represent a significant improvement over the previous version of the Galux v2 scoring function, which achieved AUROCs of 0.71, 0.50, and 0.84 for the same libraries. The method’s consistent performance across various datasets highlights its robustness and generalizability, making it a promising tool for a wide range of applications in therapeutic antibody discovery targeting various target proteins.

**Figure 2.**
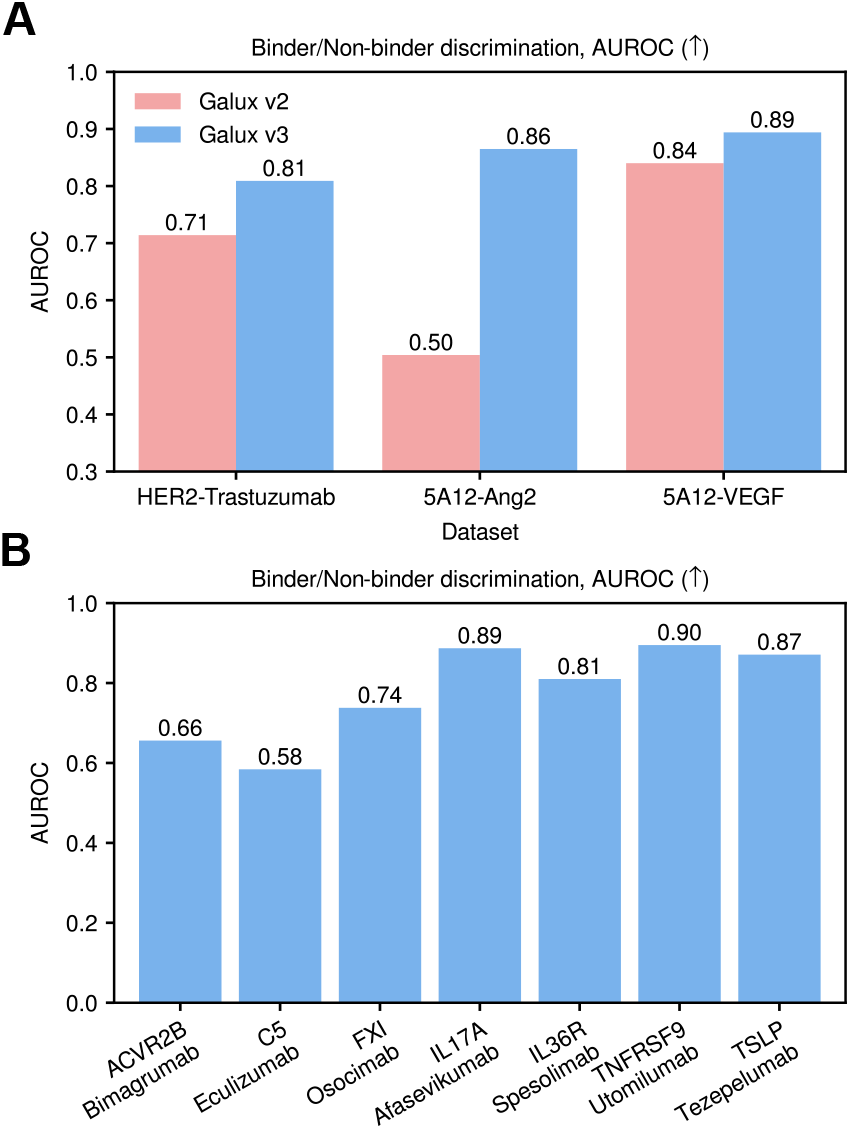
Benchmark result of binder/non-binder discrimination. (A) AUROC of Galux v2 and Galux v3 for the HER2–Trastuzumab mutant library, 5A12–VEGF, and 5A12–ANG2 mutant library (B) AUROC of Galux v3 for the IgDesign dataset

### 2.2 *In vitro* binding validation of *in silico*-designed antibodies in the scFv format across eight therapeutic targets

As illustrated in Figure 3, our design pipeline begins with the structure of a target protein (either experimentally resolved or predicted) and a designated epitope. This epitope is defined by two to five residues selected to inhibit the binding of functional partner proteins. Depending on the specific target, these residues are also strategically chosen to introduce pH dependence or subtype selectivity. Initially, we computationally generate a diverse set of antibody candidates targeting this epitope. These candidates undergo post-scoring, from which approximately 10^6^ sequences, assembled from 10^2^ light chain and 10^4^ heavy chain sequences, are selected for experimental evaluation. The selected antibodies are then displayed on yeast surfaces in the scFv format and subjected to binder screening. After three to four rounds of biopanning, enriched binders are identified.

**Figure 3.**
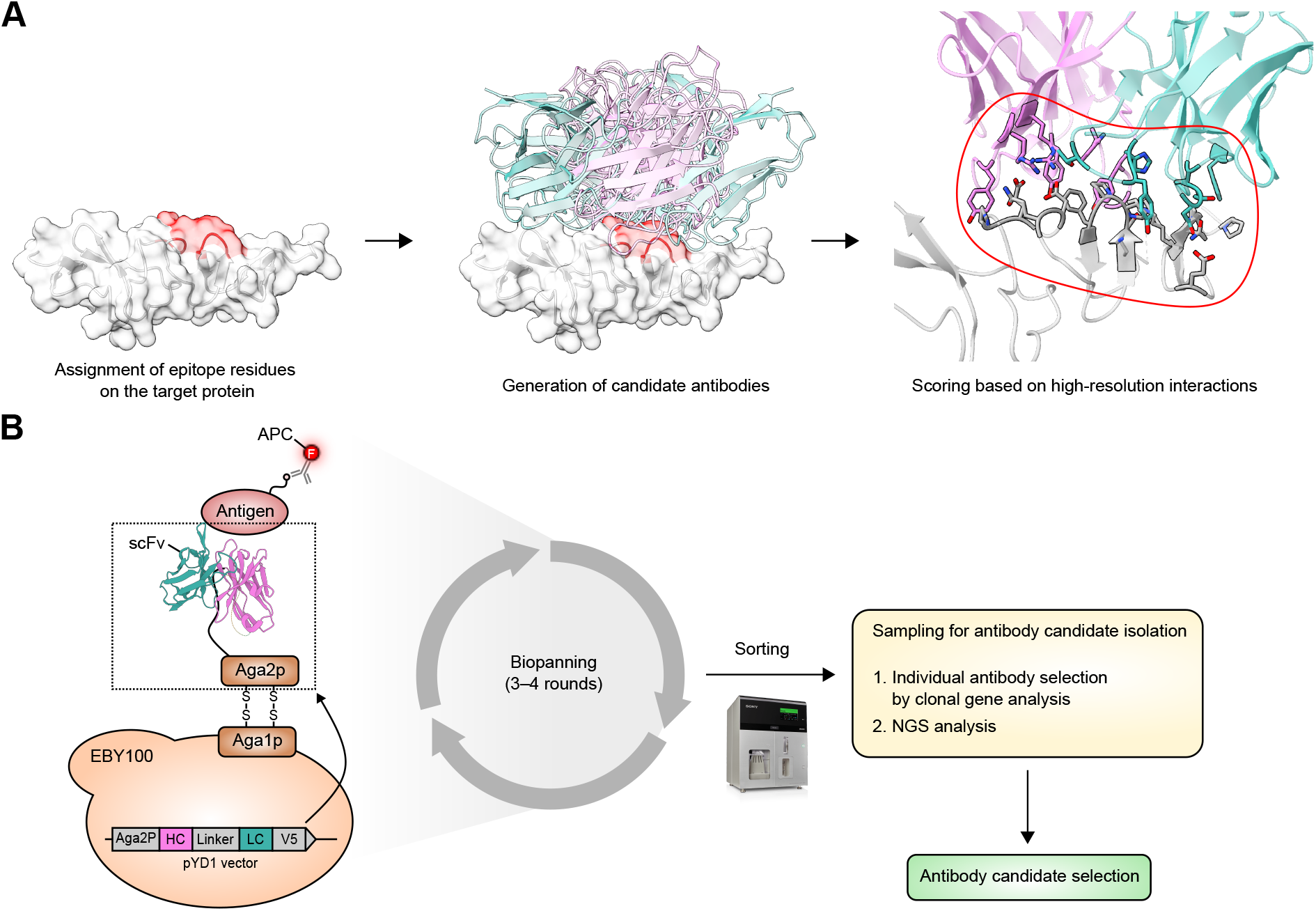
*De novo* antibody design and *in vitro* validation protocols applied to eight protein targets. (A) *In silico* epitope-specific antibody design process, including epitope assignment, antibody generation, and high-resolution scoring. (B) Yeast display-based screening process of binding antibodies.

#### Target information and design strategy

*De novo* antibody binders were designed for eight therapeutically relevant target proteins: Programmed Death-Ligand 1 (PD-L1), Human Epidermal Growth Factor Receptor 2 (HER2), S468R (or S492R when residues are numbered from the signal peptide) mutant of Epidermal Growth Factor Receptor (EGFR), Activin type 2A and/or 2B receptors (ACVR2A/B), Frizzled class receptor 7 (Fzd7), Activin receptor-like kinase 7 (ALK7), Interleukin11 (IL-11), and Cluster of Differentiation 98 heavy chain (CD98hc).

For all targets, the *de novo* design protocol was applied. For ACVR2A/B, an additional approach, termed “paratope redesign,” was employed to assess its effectiveness as an intermediate protocol between *de novo* design and CDR redesign. In paratope redesign, spatial restraints were applied to match the binding orientation observed in reference antibody–protein complexes (PDB entries 5NH3 and 5NGV [11]) instead of using epitope restraints. The CDR loops and adjacent residues were redesigned, without utilizing the reference sequence information for the redesigned region. This approach leverages structural information to guide the binding pose when such data is available. *De novo* design and paratope redesign sequences were incorporated into the library at a 7:3 ratio for screening.

#### Identification of designed antibodies with target-specific binding

Following the construction of the designed yeast library and three to four rounds of biopanning (Figure S12), multiple binder candidates were identified, and their DNA sequences were analyzed. As illustrated in Figure 4, selected binders were individually validated for specific interactions with their respective target proteins, with no detectable binding observed against multiple off-target proteins in the yeast display system (Figure S13). For ACVR2A/B, the library screening yielded binders derived from both paratope redesign and *de novo* design approaches.

**Figure 4.**
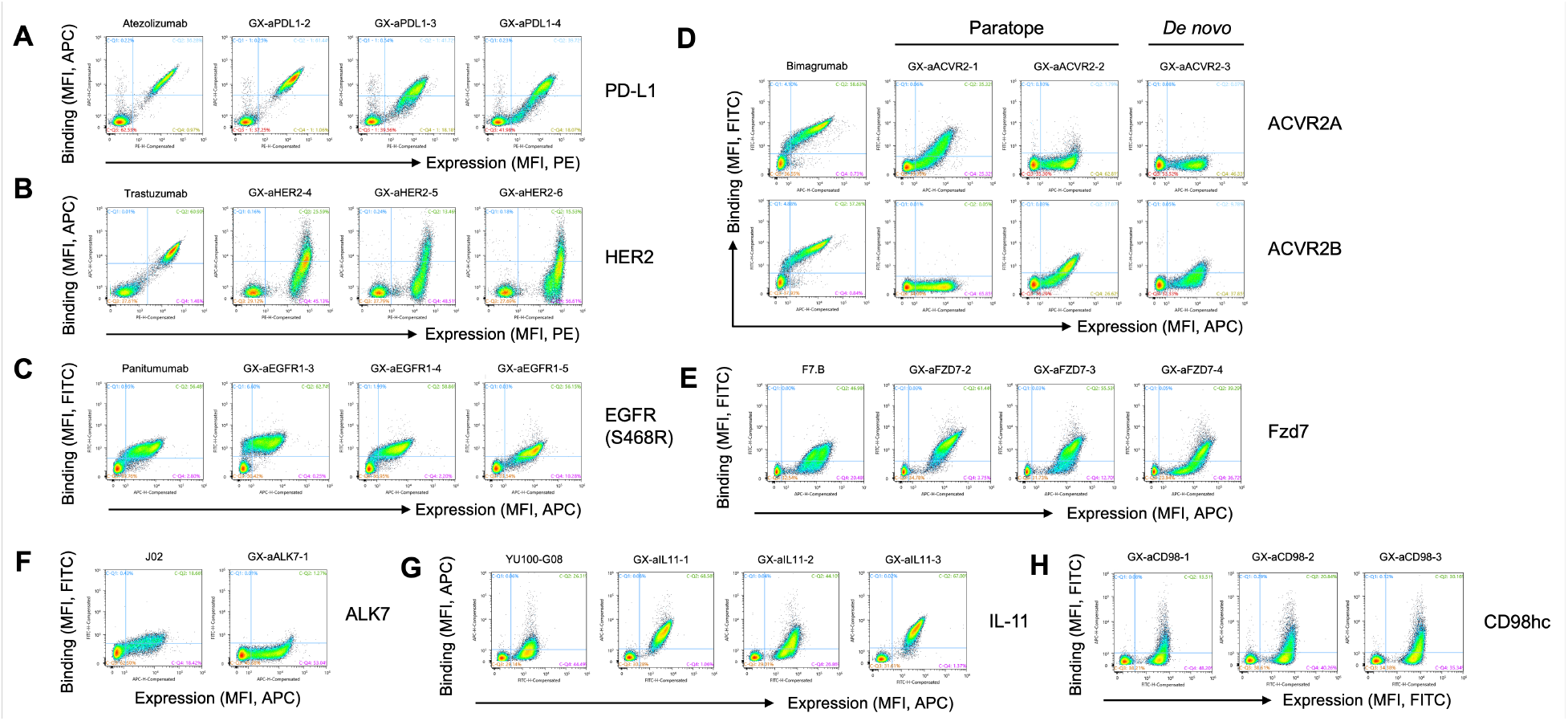
Fluorescence-activated cell sorting (FACS) data demonstrating target binding of single scFv clones using a yeast surface display system across eight targets: (A) PD-L1, (B) HER2, (C) EGFR-S468R, (D) ACVR2A or ACVR2B, (E) Fzd7, (F) ALK7, (G) IL-11, and (H) CD98hc. The prefix ‘GX-’ denotes antibodies designed by GaluxDesign and identified through screening. Reference antibodies targeting the same epitope were used for (A), (B), (C), and (D). For (E), a reference antibody targeting a different epitope was used due to the unavailability of one targeting the same epitope. For (F) and (G), a reference antibody with an unidentified epitope was used due to the unavailability of an antibody with a known epitope.

#### Epitope validation through competition assay

To verify whether the designed antibodies bind to the designated epitope regions, competition assays were performed using reference antibodies known to target the same epitopes when available (Subsubsection 4.3.3). A reduction in binding signal in the presence of the reference antibody, compared to its absence, suggests that the designed antibodies may target the intended epitope, as illustrated in Figure 5: (A) PD-L1, (B) HER2, (C) EGFR-S468R, (D) ACVR2A/ACVR2B, and (E) IL-11.

**Figure 5.**
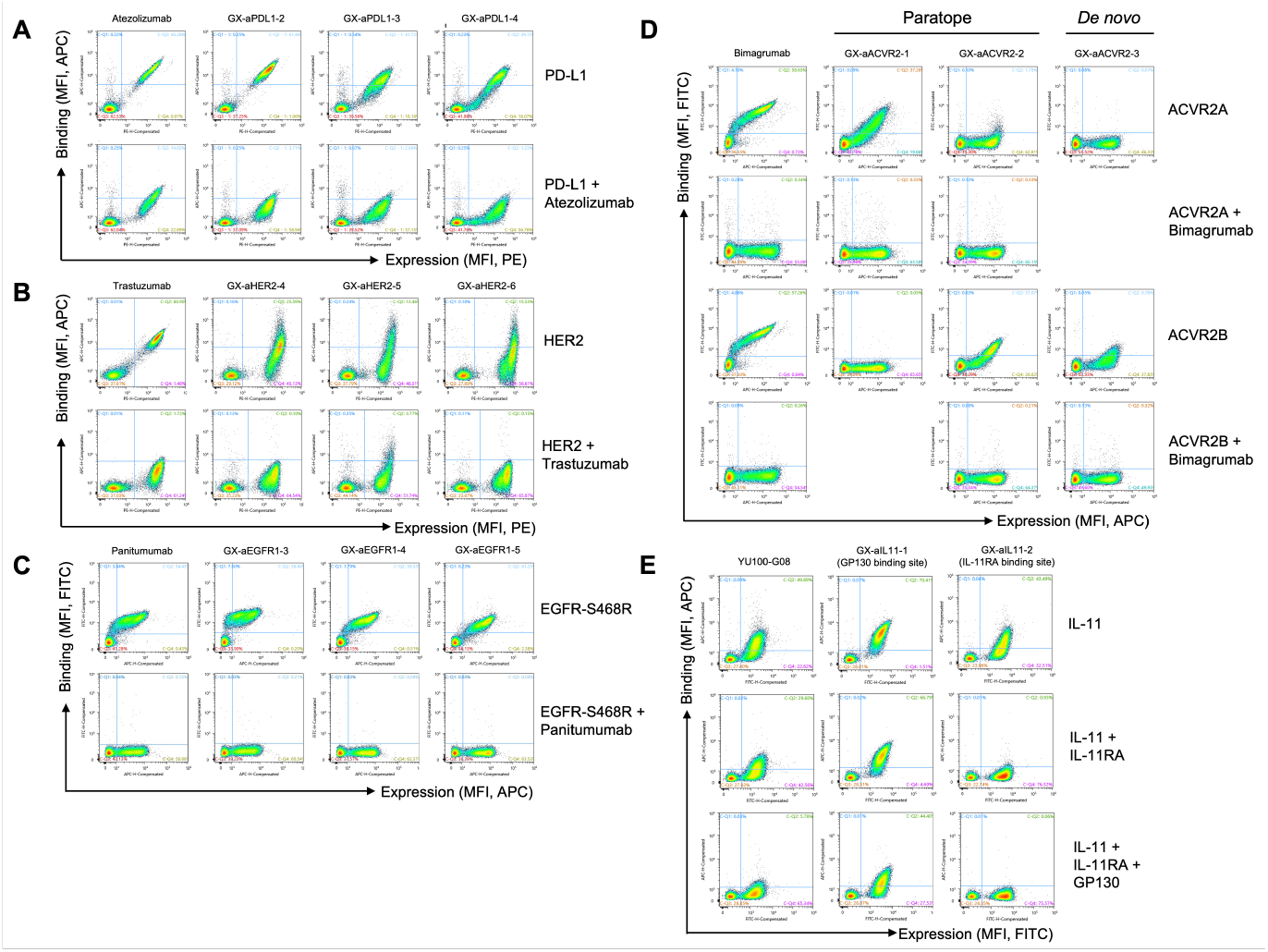
Epitope validation of selected binders through competition assays (A-D) with reference antibodies targeting the same epitope for (A) PD-L1, (B) HER2, (C) EGFR-S468R, and (D) ACVR2A or ACVR2B, and (E) with the native receptors IL-11R*α* and GP130 for IL-11.

#### Novelty and diversity of the designed antibodies

To assess the novelty of the designed antibodies, we compared their sequences with those of known antibodies targeting the same target in the PDB. Notably, a search of the RCSB PDB revealed no publicly available antibodies in complex with either IL-11 or ALK7. For other targets, most of the identified binders showed CDR-H3 sequence identity below 50% when compared to the most similar sequence in the PDB, confirming the novelty of the designed sequences. Additionally, the binders exhibited substantial sequence diversity, with an average CDR-H3 sequence identity of around 30% among those targeting the same protein.

Figure 6 highlights the novelty of the designed antibodies through two examples: an anti-Fzd7 antibody and an anti-PD-L1 antibody, both identified as target-specific binders and overlaid with their most structurally similar counterparts in the PDB. The predicted complex structures precisely target the designated sites while binding in distinct orientations from the closest antibodies in the PDB. This demonstrates that our design method can generate antibodies with novel binding poses.

**Figure 6.**
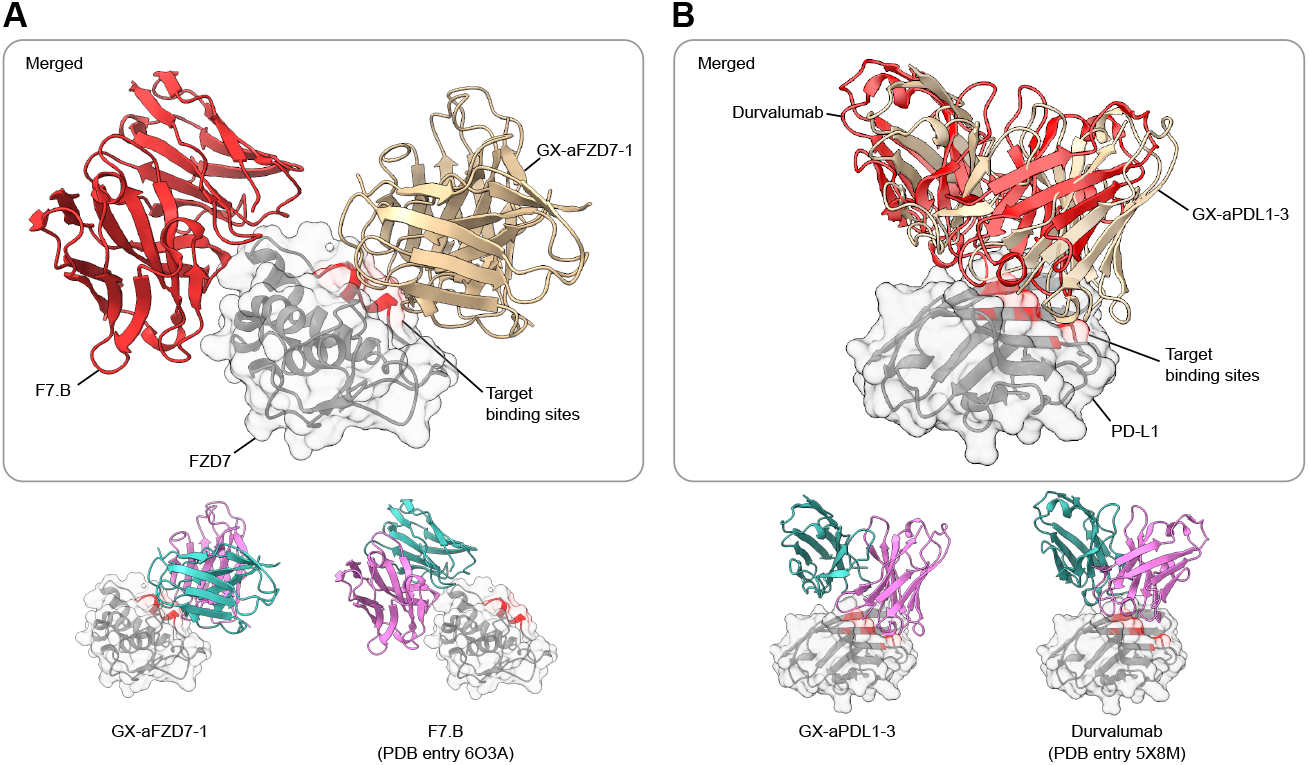
Comparison of the predicted binding poses of the designed antibodies with their most structurally similar counterparts in the PDB. (A) GX-aFZD7-1 (tan) overlaid with F7.B (red). (B) GX-aPDL1-3 (tan) overlaid with durvalumab (red).

#### Validation of designed antibodies for target subtype and mutant specificity

One of the key advantages of *de novo* antibody design over conventional discovery methods is its ability to tailor antibodies with specific binding properties. Depending on the application, cross-reactivity with related proteins may or may not be required, and both scenarios can be efficiently handled computationally.

To demonstrate this capability, two targets, EGFR-S468R and Fzd7, were selected. For EGFR-S468R, interactions were designed to distinguish between the wild-type and S468R mutant, ensuring selective binding to the mutant form (Figure 7A). Similarly, for Fzd7, antibodies were designed to bind exclusively to Fzd7, despite homologous Frizzled class receptor 1 and 5 (Fzd1 and Fzd5) sharing high sequence similarity. As shown in Figure 7B, the identified binders exhibited precise specificity for the intended subtype. These findings highlight the power of *de novo* design in achieving highly specific binding profiles, surpassing the limitations of conventional antibody discovery methods.

**Figure 7.**
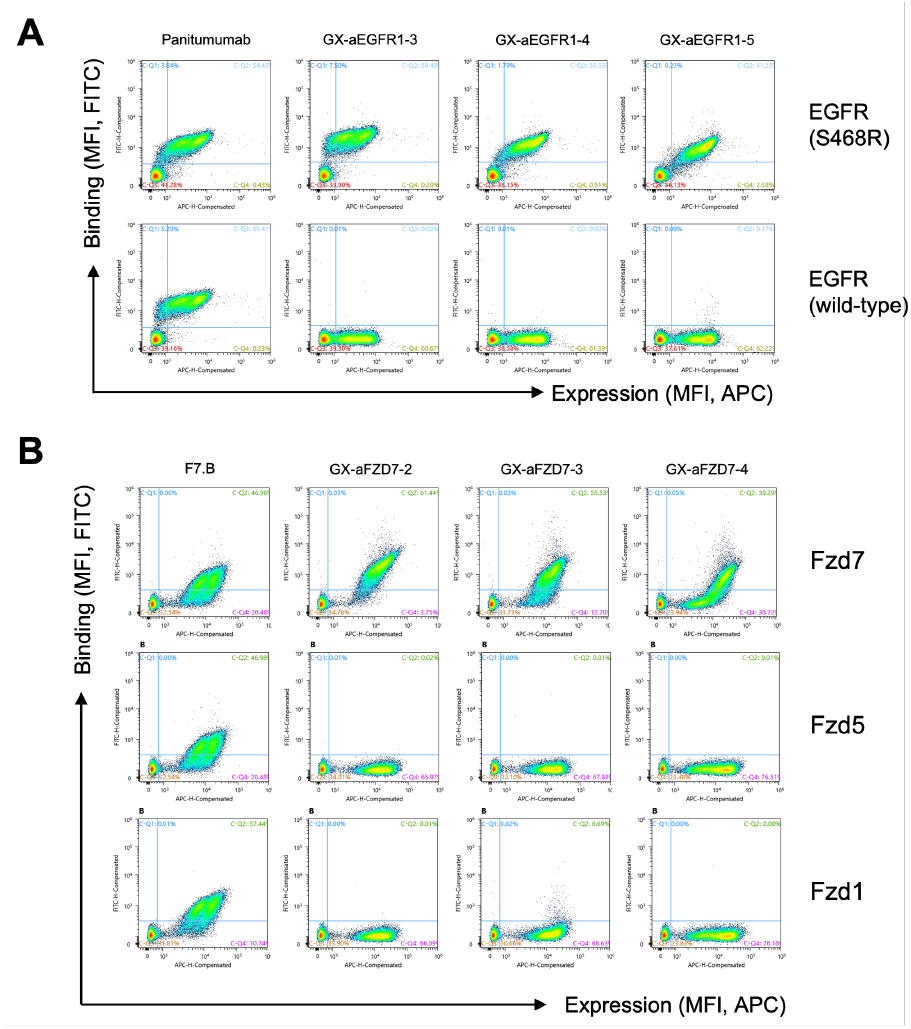
Specific binding of *de novo* designed antibodies against (A) EGFR-S468R mutant over wildtype EGFR, and (B) Fzd7 over Fzd1 and Fzd5.

Figure 8 illustrates the structural interactions formed by the designed EGFR-S468R binder, GX-aEGFR1-5, based on the predicted complex structure. Structural analysis reveals that the R468 residue plays a critical role in binding, mediated by interactions with surrounding aromatic residues and salt bridges with CDR-L3 and CDR-H3 residues. Consequently, this antibody is unable to bind to the wild-type form, in which the amino acid at position 468 is SER.

**Figure 8.**
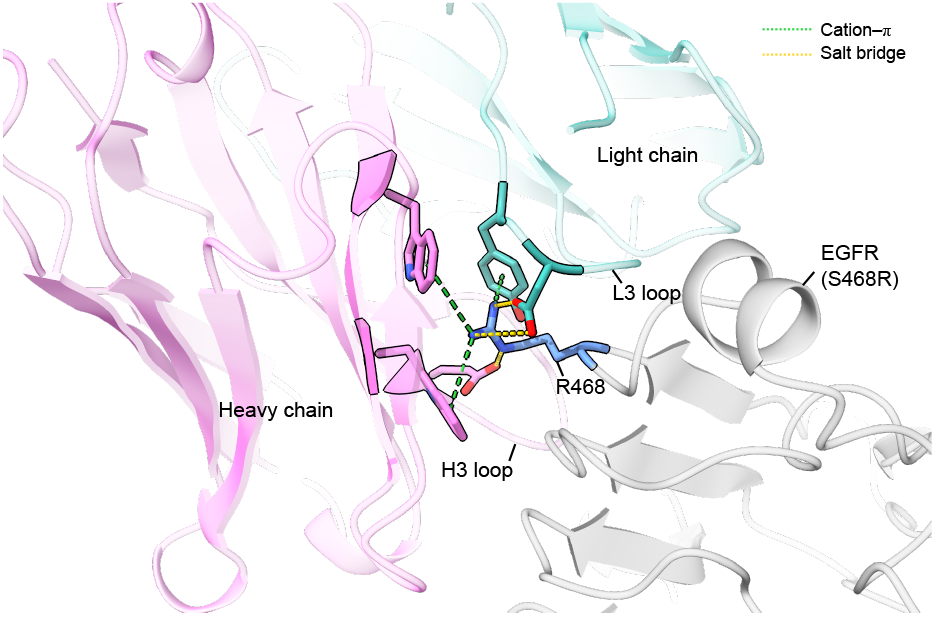
Key interactions of the EGFR-S468R binder GX-aEGFR1-5. R468 from EGFR-S468R forms salt bridges with negatively charged residues in the CDR-H3 and CDR-L3 loops and interacts with surrounding aromatic residues in the antibody.

#### *De novo* design in the absence of experimentally resolved target protein structure

ALK7 is a receptor in the Transforming Growth Factorbeta (TGF-*β*) family, primarily involved in the regulation of cell differentiation, growth, and immune responses. Thus, ALK7-targeting antibodies could offer a promising therapeutic approach for various diseases, including cancer, inflammation, and metabolic disorders.

However, *de novo* design of an anti-ALK7 antibody presents a challenge, as the ALK7 structure has not been experimentally determined. In this study, we successfully designed ALK7 binders using its predicted structure through GaluxDesign, with target-specific binding validated in Figure S13E. The target epitope residues were manually selected based on the predicted complex structure of ALK7 and its ligand, Activin E, to inhibit ALK7 function.

#### Atomic-level interaction analysis of designed antibodies with their target proteins

GaluxDesign was trained on atomic-level physicochemical interactions between proteins to guide antibody design. Analysis of the predicted complex structures further confirms the favorable formation of physicochemical interactions between the designed antibodies and their targets.

Figure 9 illustrates the interactions of the *de novo* designed antibodies GX-aPDL1-3 (anti-PDL1) and GX-aHER2-5 (anti-HER2), both identified as target-specific binders. The anti-PD-L1 antibody GX-aPDL1-3 establishes stable salt bridge interactions between ASP residue in the CDR-H1 and CDR-H3 loops and PD-L1’s R96 and R108, respectively, along with hydrophobic interactions distributed across the binding interface (Figure 9A). Similarly, the anti-HER2 antibody GX-aHER2-5 forms key interactions through hydrogen bonding between THR and TYR residues in the CDR-H1 loop and HER2’s Q561 and D560, a salt bridge between an ARG residue in the CDR-H3 loop and D560, and hydrophobic interactions around HER2’s position 572 (Figure 9B).

**Figure 9.**
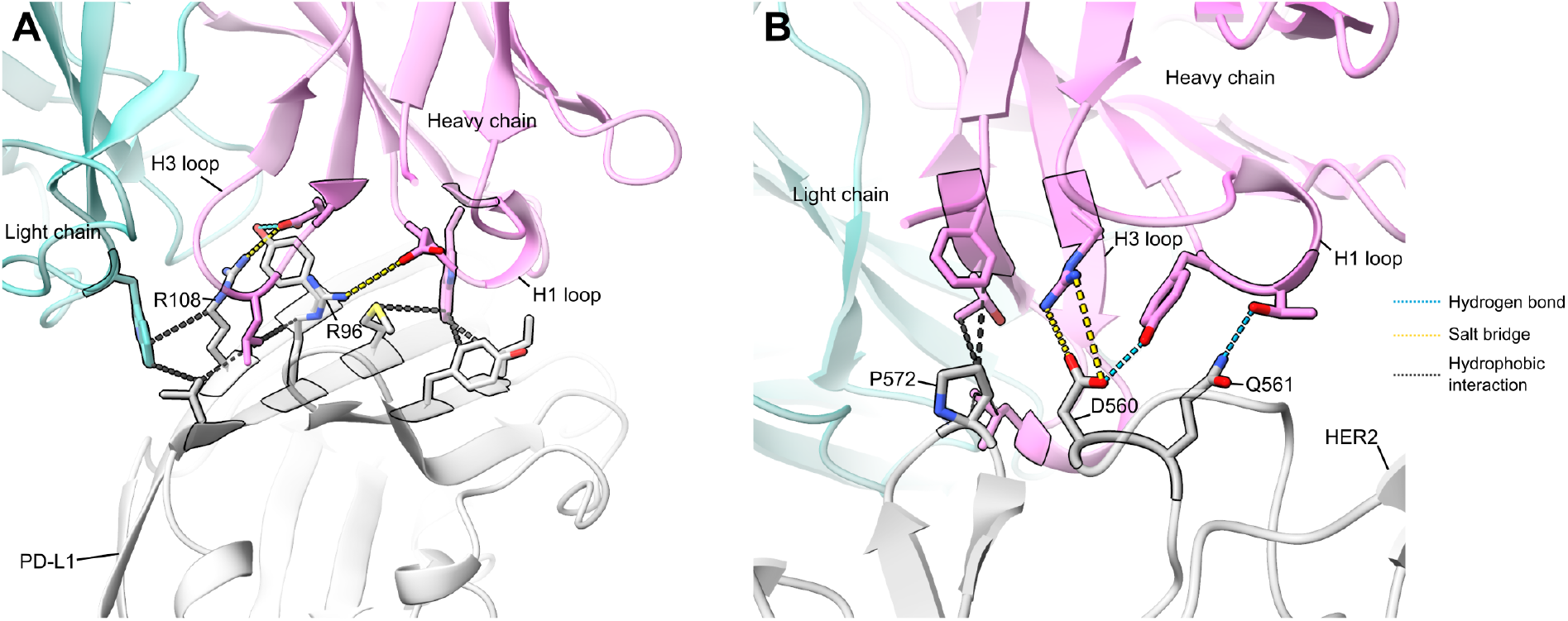
Interaction analysis of the designed antibody binders. (A) The anti-PD-L1 antibody GXaPDL1-3 forms key interactions with PD-L1, including two salt bridges between ASP in the CDR-H3 and CDR-H1 loops and the residues R108 and R96 of PD-L1, respectively, along with hydrophobic interactions distributed across the binding interface. (B) The anti-HER2 antibody GX-aHER2-5 forms key interactions through hydrogen bonding between THR and TYR in the CDR-H1 loop and the residues Q561 and D560 of HER2, a salt bridge between arginine in the CDR-H3 loop and D560, and hydrophobic interactions around P572 of HER2.

### 2.3 Cryo-EM structure determination of a designed anti-PD-L1 antibody

To experimentally verify our design, we determined the high-resolution cryo-electron microscopy (cryo-EM) structure of the PD-L1–GX-aPDL1-3 complex at 3.3 Å resolution (Figure 10A). The determined structure showed exceptional agreement with the predicted complex structure (Figure 10B), with an interface C-alpha root-mean-square deviation (iRMSD) of 1.1 Å. This close structural alignment experimentally confirms that our antibody, designed to bind a novel epitope (as shown in Figure 6B), indeed adopts the intended binding pose upon complex formation with PD-L1. Detailed analysis of the cryo-EM structure further revealed that the key binding interactions—including hydrogen bonds, salt bridges, and hydrophobic interactions—precisely match those predicted by the designed model. These results confirm the high fidelity of our computational design method in generating antibodies with precisely defined binding modes.

**Figure 10.**
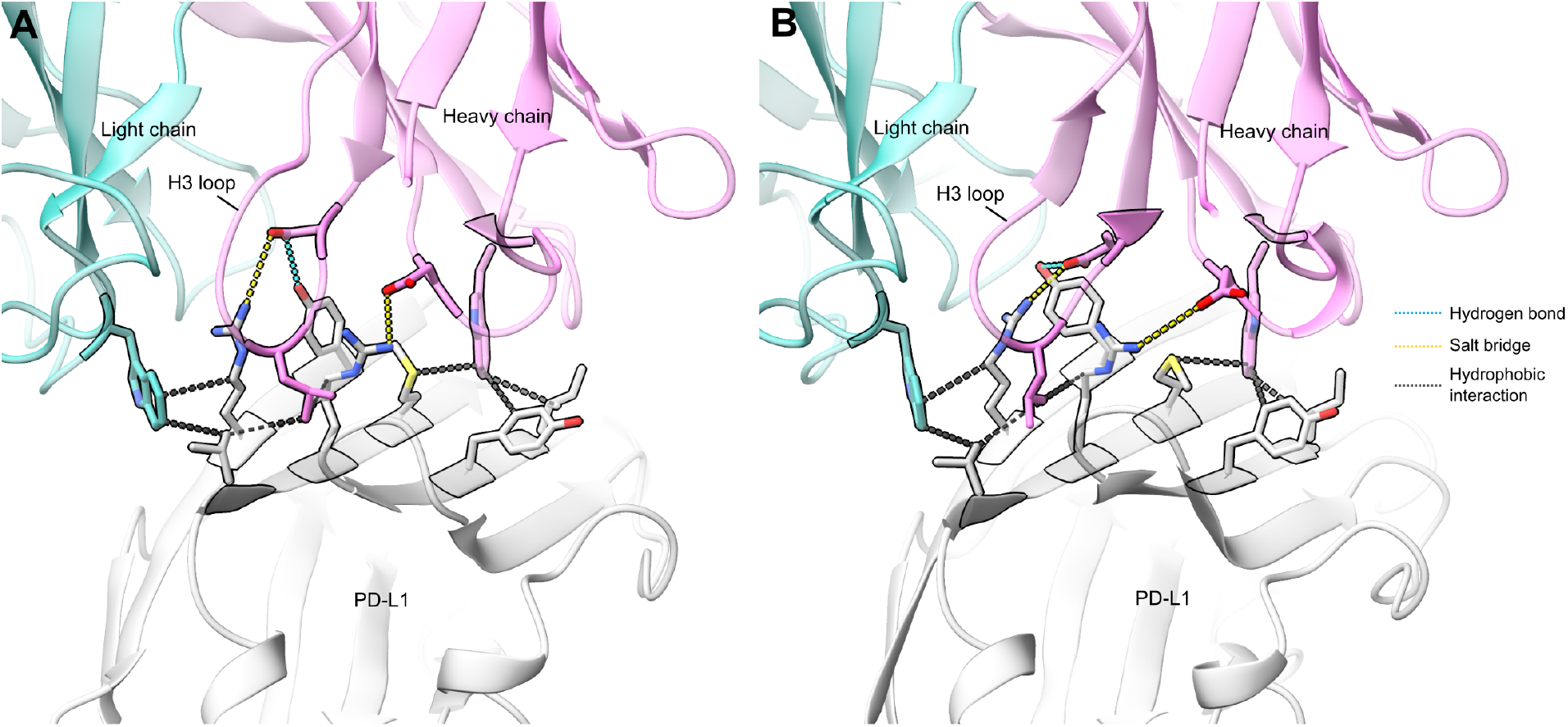
Comparison of (A) determined Cryo-EM structure (3.3 Å resolution) and (B) designed model structure of a designed anti-PD-L1 antibody (GX-aPDL1-3). The determined Cryo-EM structure, with iRMSD of 1.1 Å to the designed structure, confirms that the designed hydrogen bonds, salt bridges, and hydrophobic interactions are successfully formed, and that the intended antibody–protein binding pose is accurately realized.

### 2.4 *In vitro* evaluation of designed antibodies in the IgG format

We characterized our designed antibodies in the IgG format for five distinct targets: PD-L1, HER2, EGFR-S468R, Fzd7, and IL-11. The *de novo* designed binders, initially validated in the scFv form, were converted into the IgG format to evaluate their binding affinity, physicochemical properties, developability, and cell-based functional activity. Where available, reference antibodies were included for comparison.

#### Binding assay and epitope validation

The binding activity of the two selected anti-PD-L1 antibodies was evaluated using enzyme-linked immunosorbent assay (ELISA) and biolayer interferometry (BLI). Both demonstrated high binding affinities in the picomolar range (GX-aPDL1-2: K_d_ = 28.3 pM, GX-aPDL1-3: K_d_ = 9.0 pM), comparable to atezolizumab (K_d_ = 12.8 pM), as shown in Table 1 and Figure S14A, B. The epitope sharing of the two designed antibodies with atezolizumab was analyzed through competition assay. The binning assay using BLI confirmed that both designed antibodies compete with atezolizumab to bind to human PD-L1, implying the epitope sharing (Figure S14C).

**Table 1:** *In vitro* binding characterization profiles of designed antibodies in the IgG format.

| Antibody Index | Affinity |  | Epitope Binning Assay<br>(competition with reference antibody) |
| --- | --- | --- | --- |
|  | ELISA<br>EC<sub>50</sub> (pM) | BLI<br>K<sub>d</sub> (nM) |  |
| atezolizumab | 168 | 0.013 | (reference) |
| GX-aPDL1-2 | 41.5 | 0.028 | competition |
| GX-aPDL1-3 | 106 | 0.009 | competition |
| trastuzumab | 25.0 | 0.05 | (reference) |
| GX-aHER2-7 | 193 | 2.62 | competition |
| GX-aHER2-8 | 63.5 | 2.31 | competition |
| GX-aHER2-9 | 49.7 | 1.04 | competition |
| panitumumab | - | 1.82 | (reference) |
| GX-aEGFR1-3 | - | 1.10 | competition |
| GX-aEGFR1-5 | - | 5.62 | competition |
| GX-aEGFR1-6 | - | 1.75 | competition |
| vantictumab | 6.49 | 0.74 | - |
| GX-aFZD7-2 | 7.10 | 1.21 | - |
| GX-aFZD7-3 | - | 7.21 | - |
| GX-aIL11-1 | - | 0.37 | - |

\* - : not available

To validate the broader applicability and robust-ness of our design platform, we performed similar *in vitro* characterizations on antibodies against a diverse panel of therapeutic targets, including HER2, EGFR-S468R, Fzd7, and IL-11. The characterized antibodies successfully bound their designated epitopes, exhibiting binding affinities that spanned a range from sub-nanomolar to low nanomolar. Notably, the anti-IL-11 antibody achieved a 0.37 nM affinity (BLI; Table 1), a crucial validation of the platform’s design capabilities against targets lacking any known antibody structures in the PDB. Beyond potent affinity, the platform demonstrated the ability to generate antibodies with exceptional specificity. This precision was highlighted by the anti-EGFR-S468R antibody, which selectively bound the mutant over the wild-type protein (Figure 11A), and the anti-Fzd7 antibody, which showed no cross-reactivity with its close homologs Fzd5 and Fzd1 (Figure 11B and Table 2).

**Table 2:** Binding selectivity of a designed anti-Fzd7 antibody, confirming its specific binding to Fzd7 with no cross-reactivity with homologous Frizzled receptors.

| Antibody Index | ELISA EC<sub>50</sub> (pM) |  |  | Cell Binding Assay EC<sub>50</sub> (ng/mL) |  |  |
| --- | --- | --- | --- | --- | --- | --- |
|  | Fzd7 | Fzd5 | Fzd1 | Fzd7 | Fzd2 | Fzd1 |
| vantictumab | 6.49 | 5.03 | 395000 | 441 | 288 | 576 |
| GX-aFZD7-2 | 7.10 | n.b. | n.b. | 777 | n.b. | n.b. |

\* n.b. = not binding

**Figure 11.**
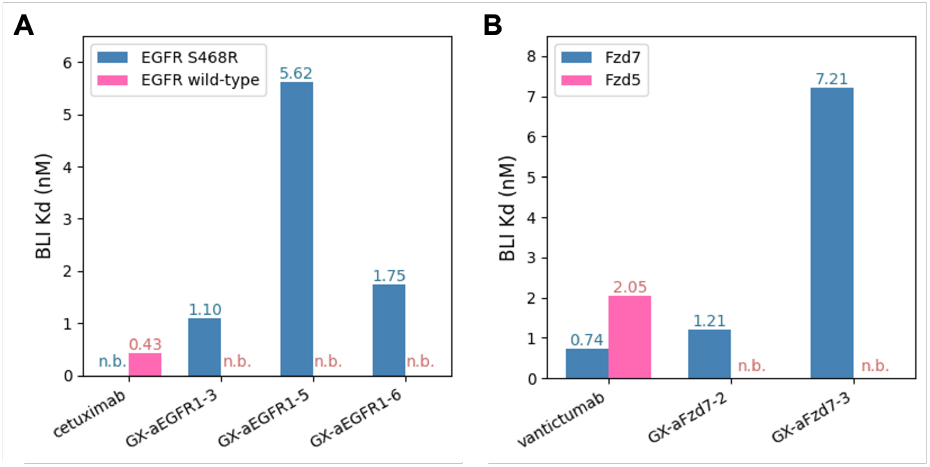
Specific binding of *de novo* designed antibodies in the IgG format, as measured by BLI assay. The antibodies show selectivity for the (A) EGFR-S468R mutant over wild-type EGFR, and (B) Fzd7 over Fzd5. n.b. = not binding

#### Developability assay

The developability of antibodies targeting PD-L1, HER2, EGFR-S468R, and Fzd7 was assessed, including productivity, thermodynamic stability, monomericity, and polyreactivity (Table 3 and Figure S15). Productivity was assessed by measuring the titer levels following transient expression of the IgG form in Expi293 cells, a mammalian expression system. The designed antibodies exhibited high productivity in the range of hundreds of mg/L, significantly exceeding the expression levels of the commercial antibody, atezolizumab. Monomericity, a critical factor influencing the antibody production process and yield, was assessed using size-exclusion high-performance liquid chromatography (SE-HPLC). The results showed that designed antibodies exhibited a very high monomer ratio. Additionally, Poly-specificity reagent (PSR) ELISA results confirmed that the antibodies lacked non-specific binding characteristics. Overall, these findings indicate that the designed antibodies targeting various targets possess stable structures with a low likelihood of aggregation and minimal non-specific interactions, which collectively highlights the generalizability of our design methodology in producing high-quality antibody candidates.

**Table 3:** Developability profiles of designed antibodies, demonstrating comparable properties to FDA-approved therapeutic antibodies.

| Antibody Index | Titer (mg/L) | Thermal Stability |  | Monomericity (%) | Polyreactivity (PSR) |
| --- | --- | --- | --- | --- | --- |
|  |  | Fab (°C) | Fc (°C) |  |  |
| atezolizumab | 84 | 64.4 | 85.4 | 96.7 | None <sup>a</sup> |
| GX-aPDL1-2 | 295 | 72.2 | 85.4 | 93.3 | None <sup>a</sup> |
| GX-aPDL1-3 | 448 | 76.1 | 84.6 | 93.5 | None <sup>a</sup> |
| trastuzumab | - | 68.3 | 83.5 | - | 2.78 <sup>b</sup> |
| GX-aHER2-7 | 538 | 68.9 | 85.1 | 100 | 2.47 <sup>b</sup> |
| GX-aHER2-8 | 617 | 69.0 | 84.6 | 100 | 4.48 <sup>b</sup> |
| GX-aHER2-9 | 685 | 69.0 | 79.4 | 100 | 4.68 <sup>b</sup> |
| panitumumab | - | - | - | - | 3.57 <sup>b</sup> |
| GX-aEGFR1-3 | 648 | - | - | 98.5 | 2.53 <sup>b</sup> |
| GX-aEGFR1-5 | 582 | - | - | 99.6 | 1.88 <sup>b</sup> |
| GX-aEGFR1-6 | 504 | - | - | 98.2 | 2.18 <sup>b</sup> |
| GX-aFZD7-2 | - | 68.1 | 89.3 | 97.6 | 1.49 <sup>b</sup> |
| GX-aFZD7-3 | - | 67.6 | 87.6 | 99.0 | 1.74 <sup>b</sup> |
| visilizumab | - | - | - | - | 7.66 <sup>b</sup> |

<sup>a</sup> Soluble membrane protein-based PSR assay
<sup>b</sup> Baculovirus-based PSR assay.
BVP Score = Test antibody signal / negative control antibody signal. (n=2)
Developability criteria: BVP Score <5.0
\* - : not available

#### Function assay

Functional evaluation was conducted using Invivogen’s PD-1/PD-L1 Blockade assay (Table 4 and Figure S16). Atezolizumab, a well-established immune checkpoint inhibitor that efficiently suppresses PD-1–PD-L1 interaction, and this function was tested in the assay. The two designed PD-L1-binding antibodies exhibited PDL1 inhibition effects comparable to those of atezolizumab.

**Table 4:** Cell-based binding assay of two designed anti-PD-L1 antibodies.

| Antibody Index | Cell Binding Assay (Raji cell, EC<sub>50</sub>) (ng/mL) | Reporter Assay (Blockade assay, IC<sub>50</sub>) (ng/mL) |
| --- | --- | --- |
| atezolizumab | 170 | 700 |
| GX-aPDL1-2 | 96 | 555 |
| GX-aPDL1-3 | 130 | 723 |

We also performed a cell-based binding affinity assay on our designed anti-Fzd7 antibody, GX-aFZD7-2. The antibody demonstrated high specificity for Fzd7, with no detectable binding to the highly homologous receptors Fzd2 and Fzd1, as shown in Table 2. This specificity profile is a key differentiator from the reference antibody vantictumab, which is known to cross-react with other Fzd homologs. Such cross-reactivity has been associated with bone-related side effects in clinical trials [12], suggesting that the exquisite specificity of GX-aFZD7-2 may offer an improved safety profile by avoiding these off-target toxicities.

In summary, our *de novo* design methodology has proven effective in generating high-quality antibody candidates for five therapeutic targets in the IgG format. Across all targets, the designed antibodies successfully demonstrated favorable biophysical properties and developability profiles. Among the key strengths of our approach is the ability to engineer fine-tuned specificity, as evidenced by the successful generation of antibodies that selectively bind to homologous targets like Fzd7 (showing no crossreactivity with Fzd5, Fzd2, and Fzd1) and EGFR (selectively binding to the S468R mutant but not the wild-type). In addition, the anti-PD-L1 antibodies, GX-aPDL1-2 and GX-aPDL1-3, demonstrated similar characteristics to the FDA-approved antibody, atezolizumab, in terms of binding activity, physicochemical properties, developability, and functional efficacy. These results highlight the effectiveness and generalizability of our design technology and its potential for direct application in therapeutic antibody discovery.

## 3 Conclusion

In this study, we successfully demonstrated the feasibility of *de novo* antibody design for targeting specific epitopes on eight distinct therapeutically relevant proteins. Unlike conventional antibody discovery methods that depend on existing immune repertoires, our approach heavily relied on computational design without incorporating prior antibody sequence/structure information. The designed anti-bodies were validated through yeast display screening in the scFv format, and the designed antibodies were further characterized in the IgG format, confirming favorable biophysical and functional properties.

Our results highlight several key achievements in computational antibody design. We successfully generated high-affinity binders with picomolar dissociation constants, antibodies capable of distinguishing subtype or mutant proteins with only a few amino acid differences, and binders targeting proteins with no known experimental structures. The high-resolution cryo-EM structure confirmed that our designed antibody successfully forms the intended binding pose, with key atomic-level binding interactions precisely matching those predicted by the designed model. These findings emphasize the strength of our structure-based molecular design strategy, which integrates atomic-level structure prediction and precision molecular design to achieve robust binding characteristics.

Beyond demonstrating the practical applicability of *de novo* antibody design, this study provides valuable insights into the advancement of computational biologics. While recent efforts in computational protein engineering have focused primarily on VHH antibody design, our work extends these capabilities to full-length scFv and IgG formats. Our findings mark a transformative milestone in antibody engineering, setting the stage for the expanded use of computationally designed therapeutics in biotechnology and pharmaceutical innovation.

## 4 Methods

### 4.1 Curation of benchmark datasets for evaluating generative model and scoring function

#### Benchmark set for evaluating *de novo* antibody generation

For *in silico* performance benchmark of *de novo* antibody design methods, we first curated a set of non-redundant antibody–protein complex structures appropriate for benchmarking purposes using the following procedure. First, antibody–protein complexes were clustered based on amino acid sequence identities. Then, we selected benchmark clusters based on the following criteria: (1) a benchmark cluster should contain at most five PDB entries and exactly one Fv cluster (identified by sequence-based clustering), and contain at least one entry where the total length of the Fv and antigen is between 300 and 1300 and whose resolution is less than or equal to 2.5 Å. Entries with non-polypeptide target proteins were discarded. Finally, for each cluster, we chose the one with the best resolution as the cluster representative, resulting in 32 structures in total.

#### Benchmark set for evaluating scoring function in binder/non-binder discrimination

Screening datasets for 10 target proteins were used to benchmark scoring methods for binder/nonbinder discrimination. One key dataset includes binding data for a HER2–Trastuzumab mutant library, as reported by Mason et al. [8]. This dataset consists of 8,955 unique binder sequences and 25,191 unique non-binder sequences, after excluding those designated as both binder and non-binder. Each sequence is identical to Trastuzumab except for variations in the CDR-H3 loop sequence. From this dataset, a subset was created by randomly sampling 268 binder sequences and 791 non-binder sequences.

Additionally, parts of the screening data for 5A12 antibody mutants [9] on CDR-H2 and CDR-L1 loops from the 5A12 mutant libraries screened against VEGF or ANG2 (5A12–VEGF and 5A12–ANG2 libraries) were used. The 5A12–VEGF library contains 642,080 sequences, with 375,267 binders, while the 5A12–ANG2 library includes 711,912 sequences, with 13,227 binders. For benchmarking, 120 binders and 1,080 non-binders from the 5A12–ANG2 dataset, and 701 binders and 499 non-binders from the 5A12–VEGF dataset were randomly selected.

The IgDesign dataset [10] which comprises libraries targeting seven therapeutic proteins was also employed. The sequences were designed based on antibody–protein complex structures, with novel CDR-H3 or HCDR (heavy chain CDR) sequences generated using IgDesign. Additionally, a portion of the sequences was generated by replacing the antibody CDR-H3 sequence with randomly selected CDR-H3 sequences from antibodies in SAbDab. In total, the dataset contains 251 binder sequences and 985 non-binder sequences.

### *4*.*2 In silico* performance evaluation of *de novo* antibody design methods

The design protocols for the two compared *de novo* antibody design methods, RFantibody and dyMEAN, are as follows. For RFantibody, we followed the instructions in the official GitHub repository (at commit e0b2188) to preprocess the antibody-protein complex structures for the benchmark into HLT format. For each protein, hotspot residues were defined as the five residues closest to the antibody, with distances measured based on C_*β*_ atom positions. Then we inferenced RFdiffusion using the author-provided weight (RFdiffusion Ab.pt) to generate antibody structure given the antibody framework sequence and hotspot residues. It should also be noted that the framework sequence provided to the model was not from hu4D5-8 (trastuzumab) as in the RFantibody preprint [3], but instead, the framework sequence of the reference antibody for each protein was used for the *in silico* test. Similarly, the CDR lengths were fixed to match those of the reference antibody for each protein. Following the default configuration of RFantibody, the number of diffusion steps was set to 50, and the noise scales were set to 1.0. After generating the antibody backbone structures, ProteinMPNN [13] was run using the script provided in the RFantibody codebase to assign appropriate amino acid identities for each CDR position.

For dyMEAN runs, we first extracted epitope definition using the script given in the official GitHub repository (at commit ecfdfa8). Heavy and light chain sequence information was prepared by masking the amino acid identities of CDR residues with hyphens. Then the dyMEAN design API was applied to obtain the designed Fv sequence and structures.

### 4.3 *De novo* antibody design and experimental validation across eight targets

#### 4.3.1 Computational design

Approximately 10^6^ target-binding antibody structures and sequences were computationally generated using the GaluxDesign v3. The key binding sites (epitopes), consisting of two to five residues for each target protein, were manually selected, prioritizing regions that could obstruct the interface of known binders. When the target structure was unavailable, predicted complex structures with known binder sequences were referenced for epitope selection. Additionally, for target proteins requiring mutant-or subtype-specific binding, distinct residues among subtypes or mutants were chosen, and surfaces that confer pH dependence were also selected.

#### 4.3.2 Yeast library construction and screening strategy for cell sorting

A DNA oligo pool from Twist Bioscience was synthesized to construct a scFv antibody. The gene was inserted into yeast plasmid (pYD1) and transformed into E. coli and EBY100 for library construction. After library construction, the yeast display screening was performed with three to four rounds of biopanning with 1 µM of the respective target protein. At each round, we gated the double positive binding region (positive for both antibody expression and target binding) and negative region (non-binding with the target) on flow cytometry and sorted the double positive population for the next round of biopanning. After finishing the library screening, NGS analysis was performed on all sorted populations from ATG Lifetech.

#### 4.3.3 Epitope binning of selected binders on yeast surface display

After each round of library screening, the sorted yeast pool was plated on an SDCAA-based agar plate to obtain yeast colonies. Individual colonies were then randomly picked up and cultured for single antibody candidate isolation. The selected antibodies were analyzed on the yeast surface to assess their binding properties with the corresponding target protein. Additionally, a competition binding assay was performed using a reference antibody to verify epitope sharing. Yeast clones were co-incubated with a fluorescent dye-labeled target protein and the reference antibody, and the binding signal was measured using fluorescence-activated cell sorting (FACS). A decrease in fluorescence intensity in the presence of the reference antibody indicated competition for the same epitope.

#### 4.3.4 Cryo-EM structure determination

GX-aPDL1-3 Fab and human PD-L1 complexes were applied to glow-discharged cryo-EM grids (Graphene film coated on Quantifoil R1.2/1.3 µm, holey carbon grids, 300 mesh Au) and plunge-frozen in liquid ethane using a Vitrobot IV (Thermo Fisher). Cryo-EM data were acquired on a Titan Krios G4 electron microscope operated at an accelerating voltage of 300 kV at the Institute of Membrane Proteins (POSTECH, Republic of Korea). All images were recorded on a Falcon 4i camera at a dose of 50 electrons per Å^2^ using EPU software (Thermo Fisher Scientific). Movies frames were gain-corrected, aligned, and dose-weighted using motion correction in RELION version 5.0 [14]. Final maps were sharpened using standard postprocessing procedures in RELION, and the resolution was estimated to 3.3 Å based on the 0.143 of the Fourier shell correlation (FSC) between two independently refined half-maps. Atomic models were built in COOT [15]. Coordinate refinement was performed in CCP-EM [16].

#### 4.3.5 IgG production and purification of the designed antibodies

##### Antibody-coding plasmid construction and IgG production

The anti-Fzd7 and anti-IL-11 antibodies were produced by Biointron (China) using the HEK293 expression system. For the remaining *de novo* designed antibodies, human codonoptimized gBlock™ (IDT) coding for the heavy chain (HC) including the VH and CH regions, and light chain (LC) including the VL and CL regions was first inserted into pcDNA™3.4-based plasmid (X11 for HC and X15 for LC) using restriction enzyme cloning. The plasmid DNA (0.5 µg) for each HC and LC dissolved in 40 µL of sterile water was then transfected into cultured Expi293F cells with ExpiFectamine™ reagent and cultured for 3 days at 37°C, 5% CO2, 125 rpm and 80% humidity to make IgG. The culture supernatant was centrifugated and filtered through a 0.45 µm filter. The titer of recombinant IgG was analyzed using BLI.

##### Antibody purification and titration

The binder antibodies were purified using MabSelect™ (Cytiva) affinity chromatography resin. Antibody concentration was determined by measuring the absorbance at A280 using Nanodrop (Thermo Fisher Scientific), and the molecular weight of the antibody was confirmed on SDS-PAGE analysis.

#### 4.3.6 Binding affinity analysis using ELISA

To determine binding affinity, an ELISA was performed for EC_50_ estimation. The optimized antigen was coated on a 96-well plate and incubated for one hour. After washing with PBS-T buffer (1 × DPBS, 0.5% Tween20), the blocking buffer (3% skim milk in PBS-T buffer) was treated for one hour. Then the antibody in a series of concentration was added to an antigen-coated plate for one hour. A secondary antibody (anti-human IgG antibody-HRP conjugation) was then added to the plate for one hour. The signal was developed by adding TMB substrate for 15 minutes, followed by the addition of a stop solution to terminate the reaction. The absorbance was measured at OD_450_.

#### 4.3.7 Binding affinity analysis using BLI

All binding assays were carried out on a Gator Prime instrument (GatorBio) using 1 × PBST (1 ×PBS, 0.05% Tween-20) as the running and wash buffer. Sensors were pre-equilibrated in buffer prior to use, and washing was performed between each assay step. The assay was conducted under optimized conditions for each target to ensure sensitivity and reproducibility.

##### PD-L1

A His-tagged PD-L1 was loaded onto Ni-NTA probes at a concentration of 6.5 µg/mL for 300 s, with the experiment utilizing Ni-NTA probes, a reference anti-PD-L1 antibody, designed antibodies, and DPBS as the buffer.

##### HER2

A biotinylated HER2 was immobilized on streptavidin (SA) sensors at 1.25 µg/mL. Association was measured by exposing the loaded sensors to trastuzumab at concentrations starting from 7.5 µg/mL in a 2-fold serial dilution series for 200 s, followed by dissociation in 1×PBST for 600 s.

##### EGFR-S468R and wild-type EGFR

The anti-EGFR-S468R antibody was immobilized on a human Fc-capture (HFc) probe at 0.6 µg/mL, and association was measured with wild-type EGFR or EGFR-S468R mutant at 3 µg/mL in a two-fold dilution series for 300 s, followed by a 300 s dissociation phase.

##### Fzd7

Fzd7 was immobilized on Ni-NTA sensors at 5 µg/mL. Association was measured by exposing the loaded sensors to the anti-Fzd7 antibody at concentrations starting from 7.5 µg/mL in a three-fold serial dilution series for 200 s, followed by dissociation in 1×PBST for 600 s.

##### Fzd5

Fzd5 was immobilized on Ni-NTA sensors at 2 µg/mL. The sensors were then incubated with antibody variants in a serial dilution series starting at 7.5 µg/mL, and association was monitored for 300 s, followed by dissociation in 1×PBST for 300 s.

##### IL-11

The anti-IL-11 antibody was captured at 2.5 µg/mL on an HFc probe, with association measured with IL-11 starting at 0.2 µg/mL in a threefold serial dilution series for 500 s, followed by dissociation for 600 s.

Raw sensor grams from all experiments were reference-subtracted and further double-referenced using either a reference channel or the lowest-concentration well of the same probe type. Kinetic parameters were obtained by global fitting to a 1:1 binding model, in which both association and dissociation phases were simultaneously fitted to derive on-rate constant (k_on_), off-rate constant (k_off_), and the equilibrium dissociation constant (K_d_=k_off_/k_on_). Normalization of loading responses was applied prior to fitting to minimize sensor-to-sensor variability. Data analysis was performed using Gator Bio analysis software to determine binding characteristics.

#### 4.3.8 Epitope binning by BLI

Epitope binding analysis was performed using BLI on a Gator Prime system (GatorBio). This assay determined whether the designed antibodies bind to the same epitope as a reference antibody, as well as to assess their binding affinity. A general competitive binding assay was performed where the antigen was immobilized first. Following baseline stabilization, the first antibody was loaded onto the antigen-coated biosensors. A subsequent association step with a second antibody was then performed to evaluate simultaneous binding. The absence of a binding response from the second antibody indicated that the two antibodies recognize overlapping epitopes or sterically hinder each other. A detectable binding response indicated they bind to distinct epitopes.

##### PD-L1

A His-tagged PD-L1 was immobilized onto Ni-NTA biosensors. In an alternative setup, either the reference antibody or the designed antibodies were immobilized first to evaluate competitive binding. For the competitive binding assay, following baseline stabilization in kinetics buffer (D-PBS – 1X, Welgene), the first antibody (reference or designed) was loaded onto the PD-L1-coated biosensors at a concentration of 250 nM for 900 seconds. A subsequent association step was performed with the second antibody at the same concentration (250 nM) for an additional 900 seconds. Dissociation was then monitored in DPBS to assess binding stability.

##### HER2

Biotinylated human HER2 protein was immobilized on streptavidin (SA) biosensors at 1.25 µg/mL. Following baseline stabilization in 1 × PBST, the primary antibody (reference trastuzumab or an engineered variant) was associated with the HER2-coated sensors at 15 µg/mL for 900 s. A second association step was then performed by introducing the alternate antibody at the same concentration for an additional 900 s, in order to evaluate whether both antibodies could simultaneously bind to the antigen.

##### EGFR-S468R

A His-tagged EGFR-S468R was immobilized onto Ni-NTA biosensors at 5 µg/mL. For the competitive binding assay, following baseline stabilization in 1 × PBST, the first antibody (either the reference panitumumab or one of the designed antibodies) was associated with the EGFR-S468R-coated biosensors at 15 µg/mL for 900 s. A subsequent association step was then performed with the second antibody, applied at the same concentration for an additional 900 s, to evaluate simultaneous binding.

#### 4.3.9 Developability assays of the designed antibodies

##### Thermal stability

To analyze the thermal stability of the binder antibody, we measured the melting temperature of the binder using a real-time PCR system. The melting temperature (T_*m*_) of the binder was determined using the Protein Thermal Shift™ Dye Kit (Thermo Fisher Scientific), which detects denatured protein levels with dye signals. The assay was performed in the following process. We prepared 125-fold diluted Protein Thermal Shift Dye, Protein Thermal Buffer, and 5 µM binder antibody, and mixed them in a PCR tube to a final volume of 20 µL. The real-time PCR device was set at 20–98.6 °C with a 0.3 °C interval, and the dye signal was measured to calculate the melting temperature.

##### Monomericity (SE-HPLC)

To evaluate the monomericity of binder antibodies, SE-HPLC was performed using a Waters Arc HPLC system (Waters). The system was equipped with a TSKgel G3000SWXL column (7.8 × 300 mm, 5 µm) (TOSOH) with a TSKgel SWXL guard column (6.0 × 40 mm, 7 µm) (TOSOH), and 2998 Photodiode Array Detector with 340–410 nm compensation. The mobile phase consisted of 0.1M Sodium Phosphate buffer (pH 7.3), with a flow rate of 1.0 mL/min and an injection volume of 50 µL. Binder samples were maintained at 10°C prior to analysis to prevent degradation. The sample concentration was 0.5 mg/mL, and the total run time was 30 minutes, with detection at 280 nm. Data acquisition and processing were carried out using the Empower 3 software. Each binder was analyzed to determine the percentage of the main peak (monomer), high-molecular-weight (HMW) aggregates, and low-molecular-weight (LMW) fragments. The relative peak areas were integrated and quantified using the Empower 3 software. The monomer peak was identified based on retention time and was compared to reference standards. The presence of HMW and LMW impurities was determined by elution profiles deviating from the expected monomeric peak.

##### Poly-specificity reagents (PSRs) assay -Soluble membrane protein-based

Soluble membrane proteins, used as PSRs, were prepared from Expi293F cells and biotinylated using EZ-Link™ Sulfo-NHS-LC-Biotin (A39257, Thermo Fisher). The PSR ELISA was then performed using a designed antibody candidate. Briefly, PSRs with optimized concentration were coated onto a 96-well plate and incubated with the antibody candidate. Streptavidin-HRP (STN-NH913, Acro biosystems) was then added, followed by the addition of TMB substrate for signal development. The reaction was stopped by adding stop solution, and the absorbance was measured at OD_450_ [17].

##### Poly-specificity reagents (PSRs) assay -Baculovirus-based

PSR assays against baculovirus particles (BVP) were performed in high-binding 96-well plates. BVP (Mednabio, E3001) was diluted 800-fold in 50 mM sodium carbonate buffer (Sigma-Aldrich, S7795, pH 9.6) and coated at 50 µL per well overnight at 4 °C. Plates were washed three times with PBS (Welgene, LB204-02; 10-fold diluted) and blocked with 5% BSA (GenDEPOT, A0100) for 1 h at room temperature. Test antibodies were applied at 1 µM for 1 h, including visilizumab (MedChemExpress, HY-P99332, positive control), trastuzumab (R&D Systems, MAB9589-100, negative control), and inhouse candidates. After six washes, bound anti bodies were detected with goat anti-human IgG Fc-HRP (Bethyl, A80-304P) at 1:4000 dilution (0.125 µg/mL) for 1 h, followed by TMB substrate for 10 min and stop solution. Absorbance was measured at 450 nm, and values were normalized by dividing each signal by the average absorbance of non-coated wells [18, 19].

#### 4.3.10 Cell-based assays of designed binders

##### Cell binding assay

Cell binding assays were performed using either Raji-APC-hPD-L1 cells (InvivoGen, #rajihpdl1), which stably overexpress human PD-L1, or HEK293A cells transiently transfected with plasmids encoding human Fzd1, Fzd2, or Fzd7 (Lipofectamine™ 3000, Thermo Fisher Scientific). Cells were incubated with serial dilutions of the designed antibodies (0–10 µg/mL) for 1 hour (at 4 °C for PD-L1 binders; at 37 °C for Fzd7 binders) in the dark. After incubation, cells were washed twice with cold FACS buffer (3% FBS in PBS) to remove unbound antibodies. For detection, either an IgG Fc-specific secondary antibody conjugated to APC (BioLegend, #410711, used for PD-L1 binders) or Goat anti-Human IgG (H+L) Cross-Adsorbed Secondary Antibody conjugated to Alexa Fluor™ 594 (Thermo Fisher Scientific, #A-11014, used for Fzd7 binders) was added at a dilution of 1:100 and incubated for 30 minutes at 4 °C in the dark. Following two additional washes with cold FACS buffer, fluorescence was measured using a NL3000 flow cytometer (NanoEnTek, #N7-0008). Data were analyzed with FlowJo™ software, and binding activity was quantified by mean fluorescence intensity (MFI) in the relevant fluorescence channel. GraphPad Prism (GraphPad Software, San Diego, CA) was used to generate binding curves and calculate the dissociation constant (K_d_) of each antibody.

##### PD-1/PD-L1 Blockade assay

Jurkat-Lucia TCR-hPD-1 effector cells and Raji-APC-hPD-L1 antigen-presenting cells were obtained from Invivogen as part of the PD-1/PD-L1 Bio-IC assay (Invivogen, #rajkt-hpd1). The cells were cultured in IMDM (Gibco) supplemented with glutamine, HEPES, and 10% heat-inactivated FBS (Corning, #35-015-CV) in T75 flasks (SPL, #70075) at 37°C in a humidified atmosphere with 5% CO_2_. The assay was performed according to the manufacturer’s instructions. Two days before the assay, cells were seeded in pre-warmed medium without selective antibiotics at a density of 5 × 10^5^ Jurkat cells/mL and 4 × 10^5^ Raji cells/mL. On the day of the assay, the cells were pelleted and resuspended in antibiotic-free medium at 2.2 × 10^6^ Jurkat cells/mL and 1.1 × 10^6^ Raji cells/mL. A 96-well flat-bottom plate (SPL, #30096) was prepared by adding 90 µL of each cell suspension per well along with 20 µL of Atezolizumab or designed binder antibodies solution at various concentrations in PBS. The plates were then incubated for 6 hours at 37°C in a humidified incubator with 5% CO_2_. 20 µL of the co-cultured cell suspension was then transferred to a 96-well white flat-bottom plate (SPL, #30196). Then, 50 µL of QUANTI-Luc 4 Lucia/Gaussia reagent (Invivogen, #rep-qlc4lg1) was added. Luminescence was immediately measured using a microplate reader (PerkinElmer, #HH35000410) with a 1,000-ms integration time. The luminescent values were initially analyzed in RLU (relative luminescence unit), This process was conducted independently on three separate plates, and the final data were the mean values obtained from these experiments.

## 5 Author Contributions

T. P., C. S., J. W., and J. Y. conceived the study and coordinated the collaborations. J. G., S. K., C. L., D. L., J. L., J. N., S. R., J. W., H. W., and J. Y. developed the predictive and generative AI methods for *de novo* protein design. Y. B., H. C., K. C., D. G., Y.-H. H., M. H., S. K., S. L., S. J. L., J. Y. M., H. O., S. O., S. P., D. H. S., M. Y. S., and M. J. Y. conducted the wet-lab validations. M. H., D. L., M. S. L., S. L., C. S., M. Y. S., J. W., and J. Y. wrote the initial draft of the manuscript. All authors contributed to the final version.

---

**Figure S12:**
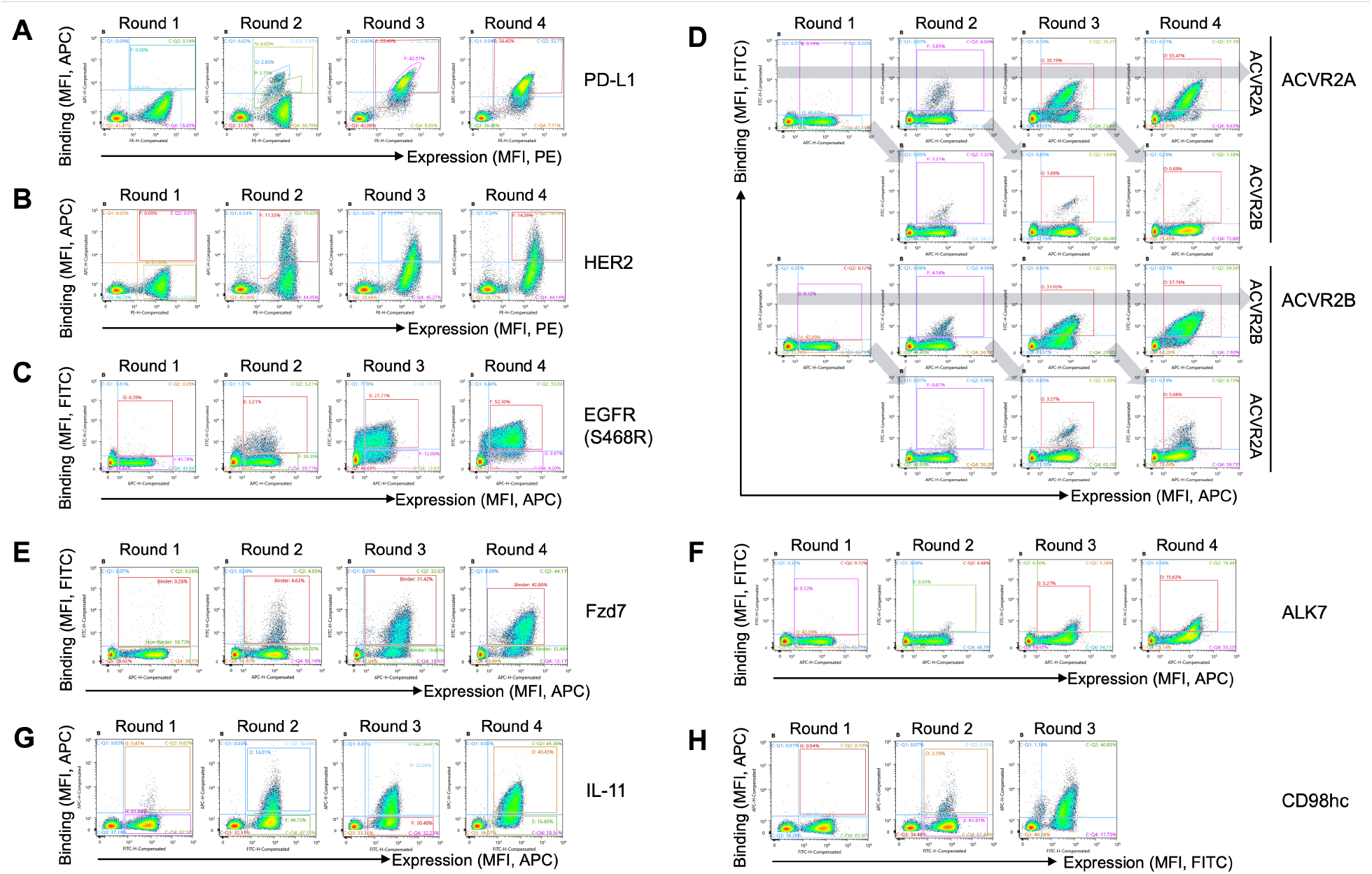
Representative FACS plots illustrating the progressive enrichment during the biopanning process for eight targets: (A) PD-L1, (B) HER2, (C) EGFR-S468R, (D) ACVR2A or ACVR2B, (E) Fzd7, (F) ALK7, (G) IL-11, and (H) CD98hc.

**Figure S13:**
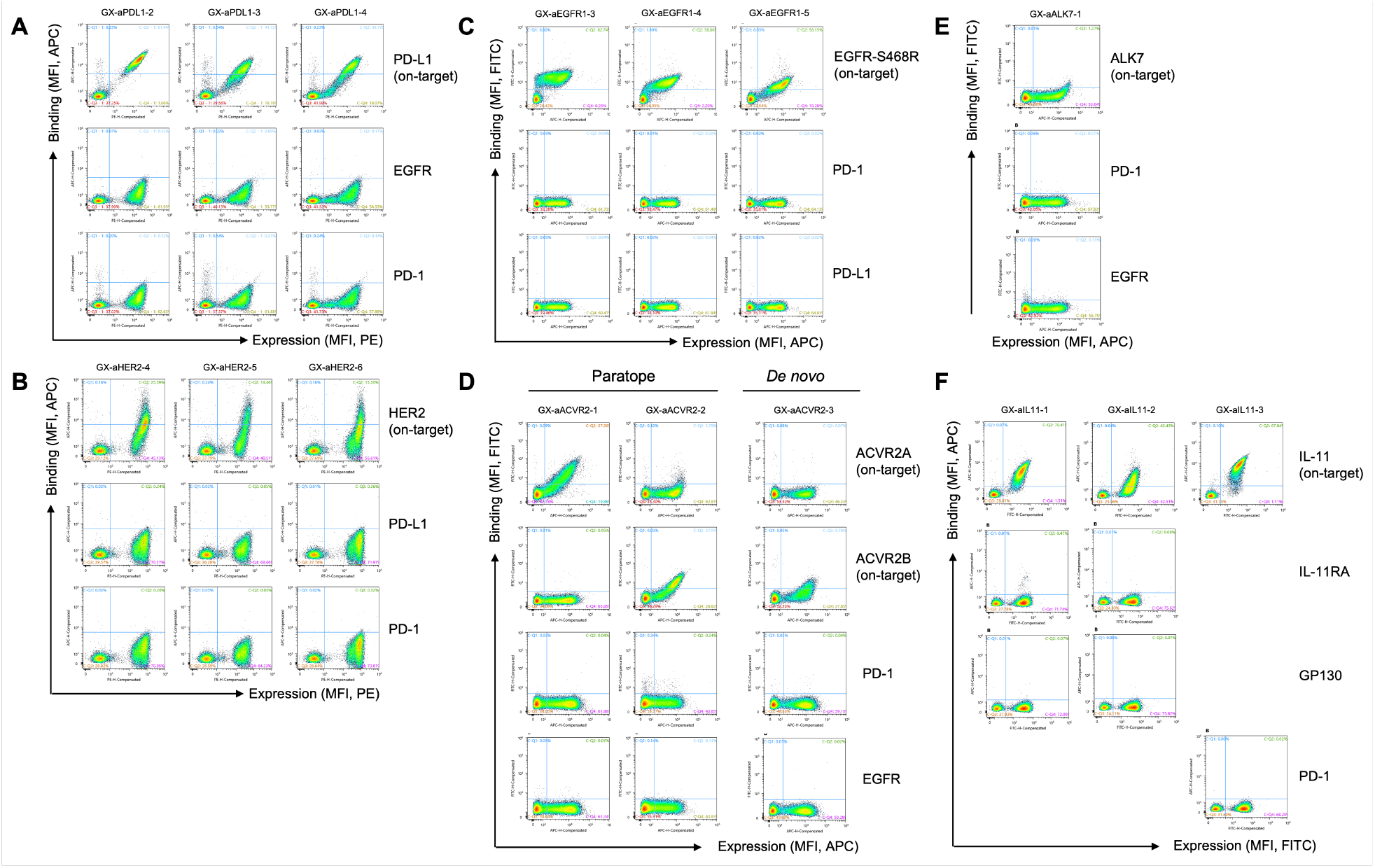
Specificity test results for the selected binder clones targeting (A) PD-L1, (B) HER2, (C) EGFR-S468R, (D) ACVR2A and ACVR2B, (E) ALK7, and (F) IL-11, assessed using a yeast surface display system.

**Figure S14:**
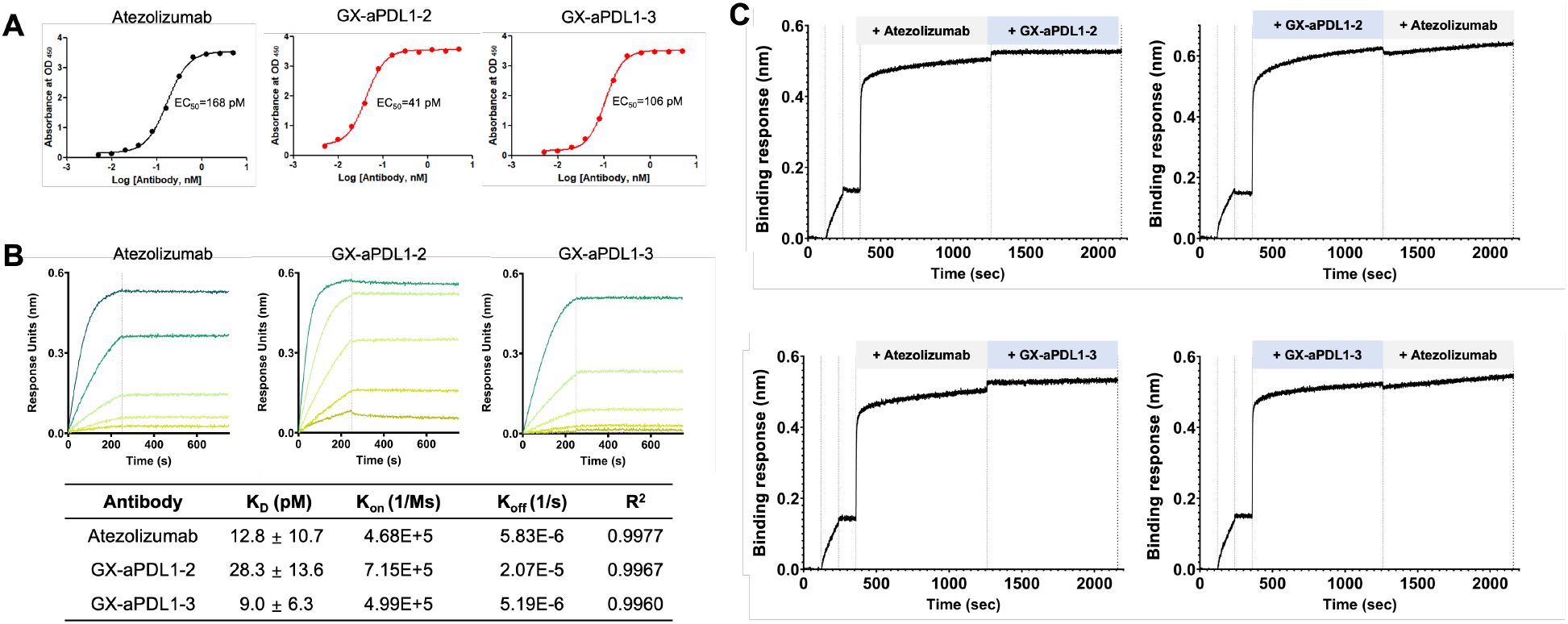
Binding affinity measurement and epitope validation of the designed anti-PD-L1 antibodies in the IgG format. (A) EC_50_ measurement using ELISA, (B) Binding kinetics analysis by BLI, and (C) Epitope binning analysis based on competition with atezolizumab.

**Figure S15:**
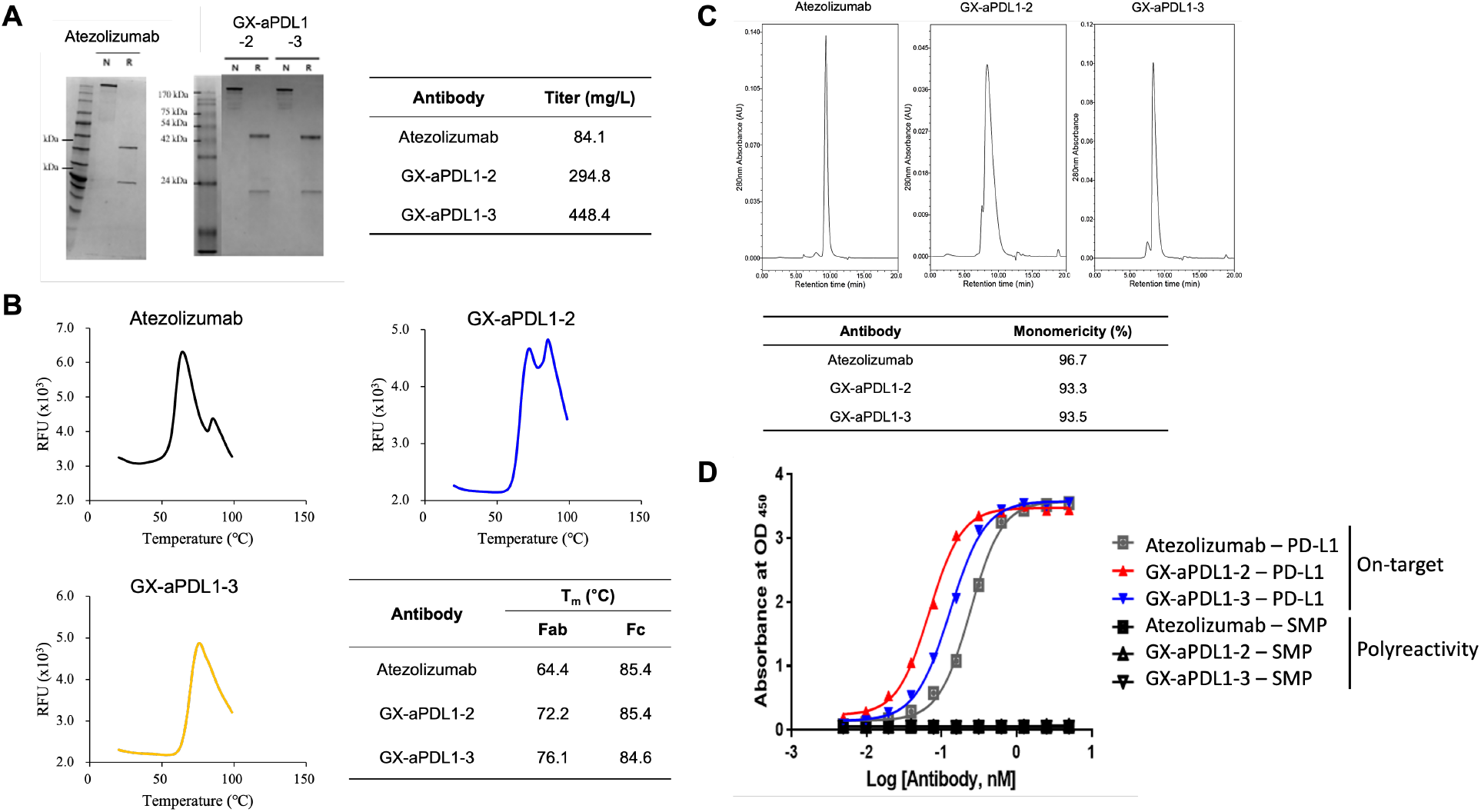
Developability assessment of the designed anti-PD-L1 antibodies in the IgG format. (A) Expression, (B) Thermal stability, (C) Monomericity, and (D) Polyreactivity.

**Figure S16:**
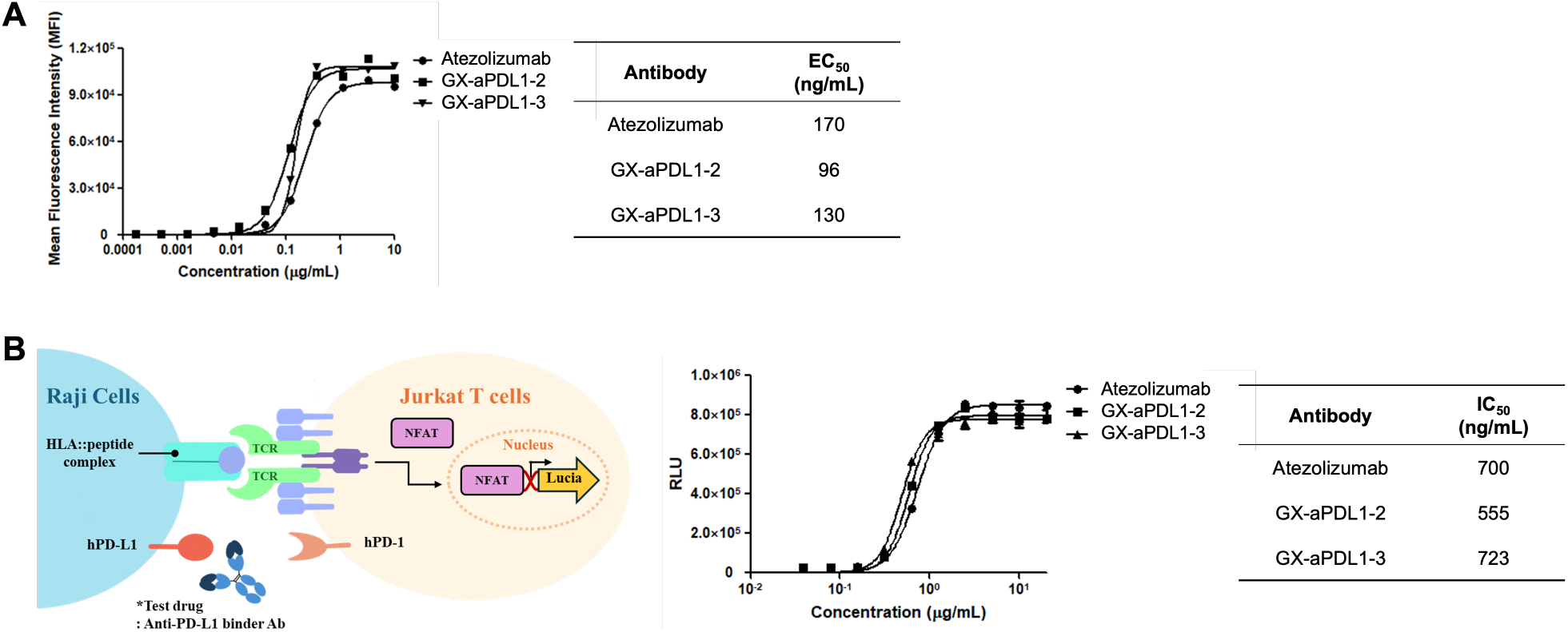
Cell-based binding and blockade assays of the designed anti-PD-L1 antibodies in the IgG format. (A) Cell-based binding assay and (B) PD-1/PD-L1 blockade assay results.

